# Molecular Heterogeneity and Early Metastatic Clone Selection in Testicular Germ Cell Cancer Development

**DOI:** 10.1101/385807

**Authors:** Lambert C.J. Dorssers, Ad J.M. Gillis, Hans Stoop, Ronald van Marion, Marleen M. Nieboer, Job van Riet, Harmen J.G. van de Werken, J. Wolter Oosterhuis, Jeroen de Ridder, Leendert H.J. Looijenga

## Abstract

**Background:** Testicular germ cell cancer (TGCC), being the most frequent malignancy in young Caucasian males, is initiated from an embryonic germ cell. This study determines intratumor heterogeneity to unravel tumor progression from initiation till metastasis.

**Methods:** In total 42 purified samples of four treatment-resistant nonseminomatous TGCC (NS) were investigated, including the precursor germ cell neoplasia in situ (GCNIS) and metastatic specimens, using whole genome- and targeted sequencing. Their evolution was reconstructed.

**Results:** Intratumor molecular heterogeneity did not correspond to the supposed primary tumor histological evolution. Metastases after systemic treatment could be derived from cancer stem cells not identified in the primary cancer. GCNIS mostly lacked the molecular marks of the primary NS and comprised dominant clones that failed to progress. A BRCA-like mutational signature was observed without evidence for direct involvement of *BRCA1* and *BRCA2* genes.

**Conclusions:** Our data strongly support the hypothesis that NS is initiated by whole genome duplication, followed by chromosome copy number alterations in the cancer stem cell population, and accumulation of low numbers of somatic mutations. These observations of heterogeneity at all stages of tumorigenesis should be considered when treating patients with GCNIS-only disease, or with clinically overt NS.

## Background

Malignant germ cell tumors of the adult testis, referred to as type II TGCTs of testicular germ cell cancer (TGCC), are the most frequent cancer in young Caucasian males (1). TGCC are thought to be initiated during early embryogenesis affecting an embryonic germ cell, and become clinically manifest during young adulthood with an annual frequency of approximately 5-12 per 100,000 men in the western world and may require “aggressive” medical treatment. These cancers are clinically and histologically classified into two variants, being seminoma (SE) and nonseminoma (NS). Both arise from a common cancer stem cell, currently referred to as Germ Cell Neoplasia In Situ (GCNIS) (2, 3), which resembles totipotent primordial germ cells (PGC) / gonocytes. Patients with proven GCNIS have a 70% chance of progression to TGCC (both SE and NS) within 7 years. SE consists of a homogeneous population of cells with similarity to GCNIS and PGC/gonocytes. About 50% of the TGCC patients present with a NS that can be composed of different histological elements, embryonal carcinoma (EC), teratoma (TE), yolk sac tumor (YST), and choriocarcinoma, either pure or mixed. The EC is the pluripotent stem cell component of NS, which can mimic normal early embryogenesis including the formation of so called embryonal bodies (EB), and thereby give rise to all differentiated components (3-5).

Although all TGCC, including mature TE, are in principle capable to metastasize, about 80-85% of the SE patients and 55-60% of the NS patients present with localized (stage I) disease. Patients with metastatic TGCC are generally cured by standard treatment regimens involving platinum compounds and additional surgery for residual TE, while only few patients show resistance to treatment (6). So far, detailed studies into the molecular profile of TGCCs and their progression stages were focused on specific genes, like *KIT* (7) and *TP53* (8-10), and on the chromosomal constitution. Analyses have revealed many changes in the (relative) number of individual chromosomes in the different tumor components (5, 11-16). Gain of the short arm of chromosome 12 is a hall mark of (invasive) TGCCs (17), but as yet no causative gene(s) have been identified. Information on driver mutations underlying the development of these NS using exome sequencing is scarce (11-14, 18-20).

In order to unravel the molecular heterogeneity of NS, we extensively investigated four rare cases of primary therapy-resistant NS and performed WGS on the primary cancer and targeted sequencing analyses on 42 enriched histological components, precursor cell populations, and metastatic lesions after treatment (Fig. 1A/B). Focus was on the early events of tumor formation, the molecular heterogeneity within the primary lesion and the retention of molecular markers in the metastatic recurrences. Additionally, data from RNA expression (RNAseq) and copy number alterations (CNA) from high throughput DNA methylation profiling and DEPArray^™^ /LowPass WGS were interrogated to decipher the evolution of the disease.

**Figure 1.**
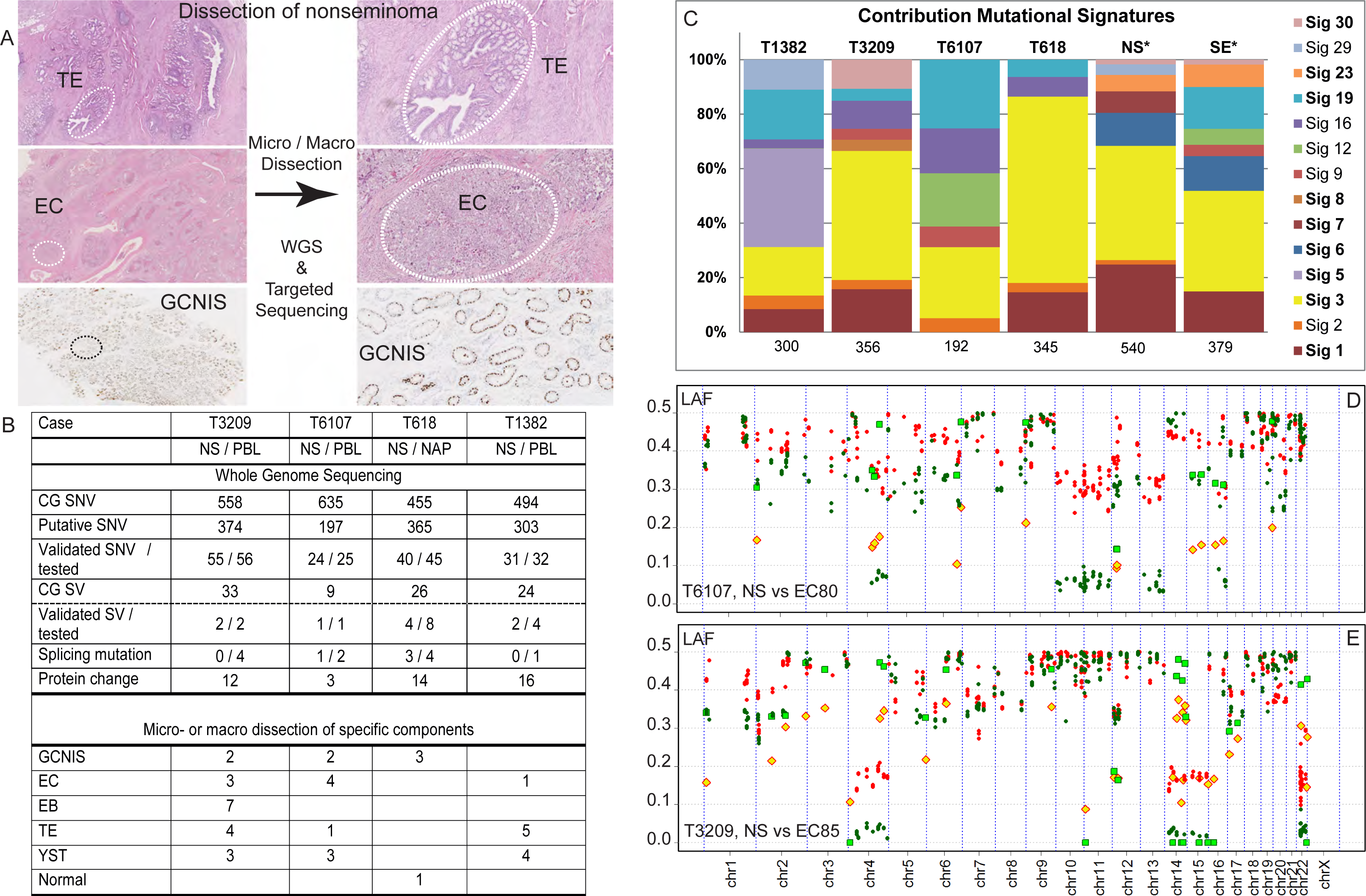
Dissection of nonseminoma. **A**. NS may consist of multiple histological elements (examples case T1382, TE and EC; case T618, GCNIS, OCT3/4 stained nuclei), which were each enriched by PALM-assisted micro dissection. The primary NS was subjected to WGS and the purified components to targeted sequencing using Ion Torrent technology. **B**. The results of the WGS are shown per case. Complete Genomics (CG) SNV were retrieved from CG output files, and putative candidates were selected after visual inspection of the reads. SNV validation was performed by mutation-specific PCR, targeted DNA sequencing and/or RNA sequencing (Supplementary Table S4). Structural variants (SV) were confirmed for selected cases using targeted sequencing. Mutations occurring near exon boundaries were evaluated for potential effect on splicing using Alamut Software. Details are provided in Supplementary Tables S4-S6. The bottom panel shows the number of purified histological components isolated and analyzed for these cases. PBL = peripheral blood leukocytes; NAP = nonmalignant adjacent parenchyma. **C**. The mutational profiles of these four NS were compared to the 30 COSMIC signatures described. The mutational signatures contributing significantly (>5%) are presented. In addition, the SNV of NS and SE cases from the Taylor-Weiner study (*) (14) were pooled for tumor subtype and similarly analyzed. The number of SNV used for the analysis is indicated below each bar. **D/E**. Profiles of lesser allele frequencies (LAF) of heterozygote SNP and relative read frequencies of SNV derived from targeted sequencing of primary NS (red) and dissected EC components (green) are shown. SNV are presented as filled symbols (NS: yellow diamonds; EC: green squares). Positions are provided on chromosomes scaled according to size.

## Materials and Methods

A brief description is provided here. Further details are provided in the Supplementary Methods.

### Patient samples

NS samples of patients with established intrinsic resistance to standard 1^st^-line chemotherapy (detailed in Supplementary Methods) were included in this study. Use of tissue samples remaining after diagnosis for scientific reasons was approved by Medical Ethical Committee of the Erasmus MC Rotterdam (The Netherlands), permission 02.981. This included the permission to use the secondary tissue without further consent. Samples were used according to the “Code for Proper Secondary Use of Human Tissue in The Netherlands” developed by the Dutch Federation of Medical Scientific Societies (Version 2002, update 2011).

### Omics analyses of patient samples

Purified tumor components (Supplementary Table S1), as defined by an experienced pathologist (JWO), were isolated from frozen tissue slices after staining for alkaline phosphatase enzyme reactivity (21), using PALM micro-dissection (Zeiss). Tumor and paired normal DNA samples were whole genome sequenced (40 times coverage) and analyzed at Complete Genomics Inc. (CG) (Mountain View, CA, USA) using NCBI build 36.3 as human reference genome and pipeline software version 2.0.2.22 (22). Lists of putative somatic DNA variants (SNV) were established from the WGS data as described in the Supplementary Methods. SNV were verified using mutation-specific Q-PCR, targeted sequencing (Supplementary Tables S2&S3), and RNA-seq. Structural variants were evaluated for gene fusions with iFuse (23). Characterization of the mutational signature was done by comparison of the trinucleotide context of each SNV to the established COSMIC signatures using the MutationalPatterns R package (v1.0) (24).

Targeted sequencing was performed by semiconductor sequencing with the Ion Torrent Personal Genome Machine with supplier’s materials and protocols (ThermoFisher Scientific) as previously described (25). Amplicons were designed to cover tumor-specific SNV, structural variants and heterozygote positions (Supplementary Table S3). Median sequencing depth was at least 250 reads. Allele frequencies were established for the heterozygous single nucleotide polymorphisms (SNP) in the matched normal samples present on the amplicons. Details for calling of SNP, SNV and structural variants in the targeted sequencing experiments are provided in the Supplementary Methods. Evolutionary trees of different samples of a specific tumor were drawn based on the LAF and SNV profiles and supported by the TargetClone tool. TargetClone was designed to reconstruct evolutionary trees for multiple samples of a cancer using allele frequencies and SNV (Nieboer MM, Dorssers LCJ, Straver R, Looijenga LHJ, de Ridder J, manuscript submitted and Supplementary Methods).

RNA samples of T6107 and T3209 were rRNA-reduced and Ion Proton sequenced (90 bases, 50 million mapped reads) using the supplier’s protocols and reagents (ThermoFisher Scientific). Generation of methylation profiles of primary tumor DNA was performed as previously described (26) or at the Microarray unit of the Genomics and Proteomics Core Facility of the German Cancer Research Center (DKFZ, Heidelberg) strictly adhering to the Illumina EPIC protocols for the T6107-YSTmeta. CNA based on methylation intensities were resolved using the Conumee package (27). DEPArray^™^ experiments on a T6107 metastatic sample, and GCNIS and YST samples of T618 were performed by Menarini Silicon Biosystems (Castel Maggiore, Italy), essentially as described (28).

## Results

### Primary nonseminoma (NS) characteristics

To address tumor heterogeneity and progression, WGS data from four primary chemo-naïve NS were exploited. Comparison of the primary tumors with the matched normal provided a set of 1239 somatic putative DNA variants (SNV) for these cases (Fig. 1B). RNAseq, mutation-specific PCR and targeted sequencing experiments validated 150 out 158 SNV and nine out of 15 structural variants (Fig 1B, Supplementary Tables S4 and S5). The identified mutations causing protein changes have been listed in Table 1. Only four SNV resulted in protein truncation and another 13 were predicted to be damaging (Supplementary Table S6). In addition, detailed information regarding structural variants, lesser allele frequencies (LAF) and chromosome copy number alterations (CNA) were obtained from the WGS and methylation profiling (details in Supplementary Figs. S1 & S2). The trinucleotide profile of single base SNV identified by WGS of the four NS was determined and compared to the established set of COSMIC mutational signatures (29). In all cases (Fig. 1C), signature 3 contributed significantly or was the predominant signature. Analysis of the pooled SNV (from whole exome sequencing) of independent cases of NS (N=18) and SE (N=18) (14), also revealed this signature to be the most prominent (Fig. 1C and further details in Supplementary Fig. S3). This signature 3 is strongly associated with mutations in *BRCA1* or *BRCA2*, genomic deletion and insertion events smaller than 100kb, and a deficiency in homologous recombination repair in breast cancer (30). In the absence of substantial numbers of indels and genomic deletion and insertion events in our WGS data, additional support for recombination repair deficiency is lacking. Furthermore, pathogenic somatic mutations in *BRCA1* or *BRCA2* were not observed in the four included TGCC, although case T1382 carried a predicted, non-pathogenic BRCA1 missense variant (Supplementary Table S6), nor in the cases of the Taylor-Weiner study (14). In addition, promoter hypomethylation and RNAseq reads observed for both genes did not support loss of BRCA function (Supplementary Fig. S4).

**Table 1.** Somatic mutation derived protein variants. Somatic mutation of genes leading to (putative) protein changes per case. The mutation type, RNA nucleotide (NT) and amino acid (AA) changes are indicated. The last three columns provide the occurrence of the DNA mutation (samples with mutation / samples successfully sequenced) in the different enriched sample types (GCNIS, histologies of the primary tumor, and metastases). n.d. = not detected, indicating that the sequencing of the specific target was not successful.

| Tumor | Symbol | ACC | Mutation Type | NT | AA | Presence of SNV in |  |  |
| --- | --- | --- | --- | --- | --- | --- | --- | --- |
|  |  |  |  |  |  | GCNIS | Histology | Meta |
| T1382 | EIF5B | NM_015904 | missense | c.541A>C | p.N181H |  | 1/1 | 7/7 |
|  | TTN | NM_003319 | missense | c.5558T>A | p.I1853N |  | 2/2 | 5/8 |
|  | LPCAT1 | NM_024830 | missense | c.1129G>A | p.E377K |  | 2/2 | 7/7 |
|  | LPCAT1 | NM_024830 | missense | c.517G>C | p.V173L |  | 2/2 | 8/8 |
|  | CHRNA2 | NM_000742 | missense | c.688G>A | p.E230K |  | 2/2 | 1/8 |
|  | KAT6A | NM_006766 | missense | c.158T>C | p.L53S |  | 2/2 | 4/4 |
|  | SYT7 | NM_001252065 | missense | c.655G>A | p.G219S |  | 1/1 | n.d. |
|  | BICD1 | NM_001003398 | missense | c.490C>T | p.R164W |  | 2/2 | 1/8 |
|  | FREM2 | NM_207361 | missense | c.7882C>T | p.R2628W |  | 2/2 | 5/6 |
|  | POLR3K | NM_016310 | stopgain | c.49G>T | p.G17X |  | 2/2 | 8/8 |
|  | SNX20 | NM_182854 | missense | c.911T>A | p.I304N |  | 2/2 | 8/8 |
|  | SETD6 | NM_024860 | missense | c.1318A>G | p.I440V |  | 2/2 | 0/6 |
|  | CA5A | NM_001739 | missense | c.721G>A | p.E241K |  | 2/2 | 8/8 |
|  | BRCA1 | NM_007300 | missense | c.745A>G | p.T249A |  | 2/2 | 4/7 |
|  | TMEM147 | NM_032635 | missense | c.104G>A | p.C35Y |  | 2/2 | 1/1 |
|  | SLCO4A1 | NM_016354 | missense | c.175C>T | p.L59F |  | 2/2 | 8/8 |
| T3209 | MTOR | NM_004958 | missense | c.3011G>C | p.C1004S | 0/1 | 13/13 |  |
|  | NRXN1 | NM_001135659 | missense | c.634G>T | p.G212C | n.d. | 6/6 |  |
|  | TAF7 | NM_005642 | nonfrs del | c.738_740del | p.I247del |  |  |  |
|  | PDLIM7 | NM_005451 | missense | c.422C>T | p.P141L | n.d. | 11/12 |  |
|  | CCKBR | NM_176875 | missense | c.1240C>G | p.R414G | n.d. | 2/9 |  |
|  | RNF219 | NM_024546 | missense | c.1940A>T | p.Q647L | n.d. | 5/5 |  |
|  | GCOM1 | NM_001285900 | missense | c.1027G>C | p.E343Q | 0/1 | 3/13 |  |
|  | ABCC3 | NM_003786 | acceptor | c.2415-5C>T | Unknown | 0/1 | 13/13 |  |
|  | ZNFX1 | NM_021035 | frs del | c.3592_3593del | p.S1198AfsX71 |  |  |  |
|  | SHOX | NM_006883 | missense | c.310G>A | p.V104M | n.d. | 8/8 |  |
| T6107 | CPSF3 | NM_016207 | missense | c.762T>G | p.D254E | 0/2 | 6/6 | 0/1 |
|  | UACA | NM_018003 | missense | c.2272G>C | p.D758H | n.d. | 3/3 | 0/1 |
|  | DNAJB1 | NM_006145 | missense | c.200A>G | p.Y67C |  |  |  |
| T618 | CROCC | NM_014675 | nonfrs sub | c.244_246 | p.Q82del | 3/3 |  |  |
|  | ARHGEF10L | NM_018125 | acceptor | c.-43-3C>T | Unknown | 0/3 |  |  |
|  | MYOM3 | NM_152372 | nonfrs sub | c.3688_3689delinsAG | p.Q1230R | 0/3 |  |  |
|  | TTF2 | NM_003594 | missense | c.634C>A | p.H212N | 0/1 |  |  |
|  | KIAA0226 | NM_014687 | missense | c.1963C>T | p.H655Y | 3/3 |  |  |
|  | SMARCAD1 | NM_001254949 | missense | c.226A>G | p.N76D |  |  |  |
|  | ASAP1 | NM_001247996 | missense | c.933G>C | p.Q311H | 0/2 |  |  |
|  | DNAJB5 | NM_001135005 | frs sub | c.1077_1093G | p.Nfs?? | 0/3 |  |  |
|  | ZDHHC6 | NM_022494 | missense | c.950G>A | p.R317H |  |  |  |
|  | CLCF1 | NM_001166212 | missense | c.468A>T | p.E156D | 0/3 |  |  |
|  | NDUFV1 | NM_007103 | acceptor | c.511-4G>A | Unknown | 1/2 |  |  |
|  | LRRC10 | NM_201550 | missense | c.125G>A | p.R42H | 0/3 |  |  |
|  | PIAS1 | NM_016166 | missense | c.1712A>C | p.D571A | 0/3 |  |  |
|  | KEAP1 | NM_203500 | missense | c.610C>T | p.R204W | 3/3 |  |  |
|  | CSRP2BP | NM_020536 | stopgain | c.412G>T | p.E138X | 1/2 |  |  |
|  | CSE1L | NM_001316 | donor | c.1619+4A>T | unknown | 3/3 |  |  |
|  | APOL3 | NM_145640 | missense | c.1066C>G | p.L356V | 0/3 |  |  |
|  | CFP | NM_002621 | missense | c.1285G>T | p.V429L | 3/3 |  |  |

### Molecular heterogeneity and evolution

For the study of the molecular heterogeneity within these histologically complex primary NS (containing EC, EB, TE and YST components, Fig 1A), matched metastases and precursor lesions, DNA was prepared from various micro- and macro-dissected components (N=42, Fig. 1A/B & Supplementary Table S1). The histological identity was determined by an experienced pathologist, and using direct alkaline phosphatase-staining for EC, EB and GCNIS in frozen tissue (examples in Supplementary Fig. S5) (21). To evaluate the allelic imbalances and the presence of SNV in these enriched specimens, amplicons were designed across the genome containing a tumor-specific SNV and additional heterozygous SNPs (Supplementary Fig. S1 and Supplementary Table S3). For the primary tumor DNA samples, an excellent agreement between the LAF profiles of the WGS and targeted analyses was observed (Supplementary Fig. S6). Analyses of the enriched samples were focused on the LAF of germ line heterozygote SNPs, the read frequencies of the SNV and the presence of specific breakpoints. Furthermore, evolutionary trees based on the LAF and the presence of SNV, and supported by TargetClone, were generated for each case. Results of these analyses will be discussed per case below.

T6107: The majority of the allelic imbalances and SNV found in the primary cancer were present and more easily detected in the enriched malignant histologies due to their increased purity and the removal of contaminating normal cells (example in Fig. 1D). Loss of heterozygosity (LOH) was clearly resolved for chromosome (arm) 4q, 10, 11, 13 and 16q for the purified EC component. The false color plot showed extensive overlap in regional allelic imbalances and SNV amongst the enriched histological components of T6107 and compared to the primary NS (Fig. 2A/B and Supplementary Fig. S7A). LOH on the chromosomes mentioned above was preserved in all histological elements (homogeneously red colored blocks indicating LAF<0.1, Fig. 2A). Sample EC21 displayed additional LOH resulting from copy losses of chromosomes arms 9q and 22q. In the GCNIS preparations (CIS30&FCIS31, Fig. 2A), very little overlap in LAF patterns with the primary NS was observed and the majority of SNV were completely absent (Fig. 2B). The yolk sack tumor metastasis in the lung (YSTmeta) showed minor overlap with the primary NS and its histological components with regard to LAF pattern and presence of SNV. The shared LOH of chromosome arm 22q between EC21 and this metastasis represented independent events based on the different parental alleles retained (Supplementary Fig. S8). Presence of chromosome arm 12p gain (Supplementary Fig. S9), and two SNV (Fig. 2B) in this metastasis demonstrated a shared origin with the primary NS. The copy number profiles of this lung metastasis and a prior retroperitoneal lymph node metastasis with the histology of mature TE displayed many novel alterations, including amplification of the MDM2 region (details in Supplementary Fig. S9). Immunohistochemistry (IHC) and fluorescent in situ hybridization (FISH) analysis confirmed the amplification of MDM2, and targeted sequencing did not reveal TP53 mutation in the lung metastasis (not shown). An evolutionary tree for this case (Fig. 3) was based on the general profiles of the LAF and SNV (Fig. 2A/C), and required two unidentified EC precursors (ECx1 & ECx2) to explain the variance between the primary NS components and the YST lung metastasis. EC21 represented a separate progression line with additional chromosome losses. Furthermore, the early ECx1 precursor containing few aberrations was the founder of the late appearing lung metastasis (YSTmeta) and likely of the mature TE in the lymph node metastasis (Supplementary Fig. S9), which both lacked many of genomic marks of the primary tumor.

**Figure 2.**
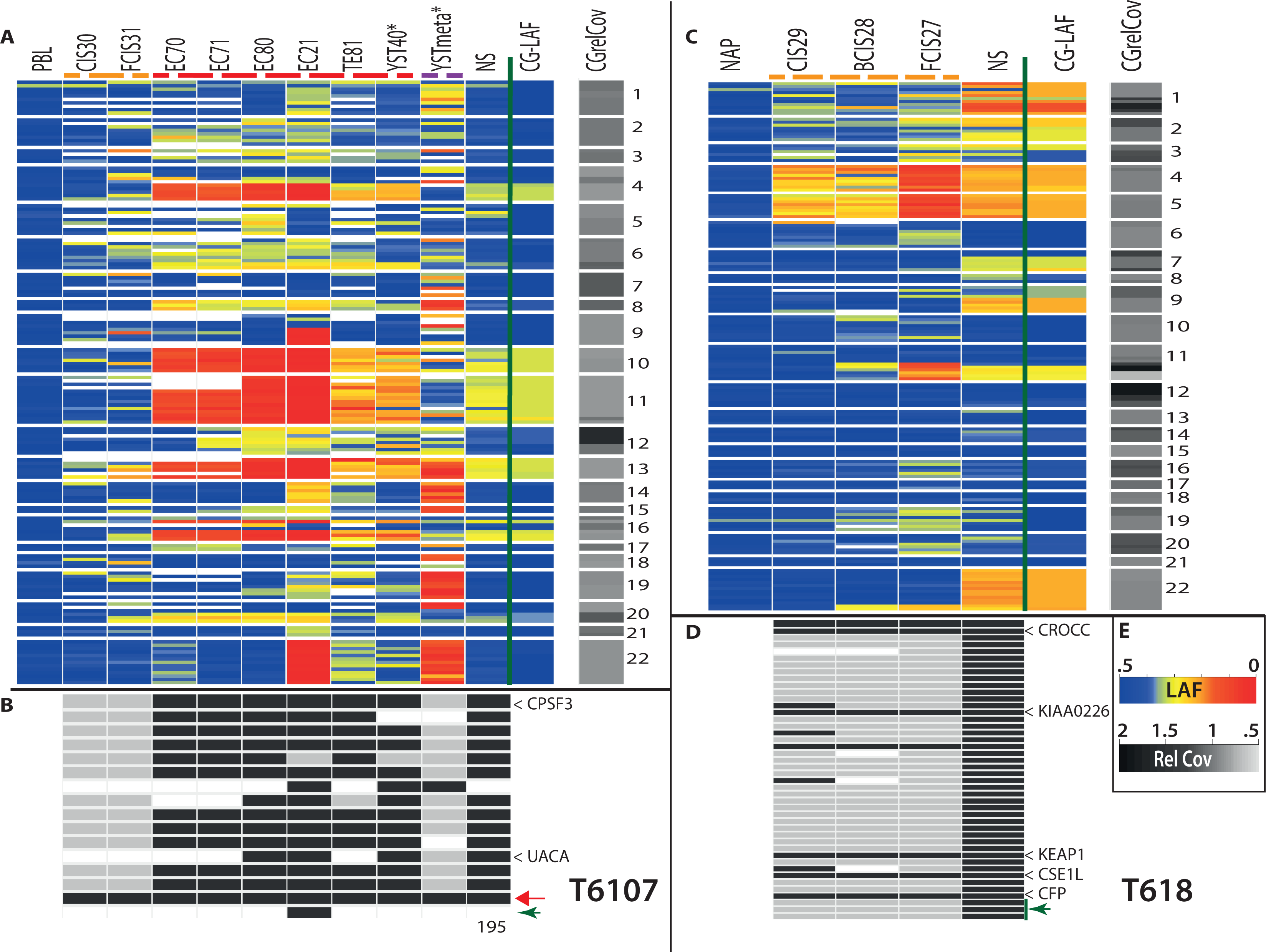
False color plots of allelic imbalances, SNV and structural variants in primary TGCC (NS) and various purified tumor components of two cases (T6107 (panels A/B) and T618 (panels C/D)). Data from (tumor) samples analyzed more than once were averaged. **Top panels A and C:** Each line represents the amplicon averaged LAF of heterozygote SNPs using the LAF color scheme in panel E (LAF). Blue indicates heterozygosity and red refers to LOH. Ordering is based on chromosomal position (indicated on the right). For comparison, the 100kb interval WGS LAF (CG-LAF) is also shown. In addition, the WGS relative read coverage (CGrelCov) data of the primary NS are provided for the specific chromosomal regions using the color scheme (Rel Cov) in panel E. Missing data are white. Sample types have been marked by colored dashed lines (GCNIS: orange, histological components: red and metastases: purple). (*) FFPE tissue blocks derived DNA samples. **Bottom panels B and D:** Occurrence of tumor-specific SNV and structural variants in the different tumor samples (grey indicates absence, black indicates more than 3% of the reads carrying the variant, missing data in white). Tumor-specific structural variants are indicated with green arrow heads. Genes with a mutation resulting in amino acid change (T618: only those observed in GCNIS) have been indicated (see Table 1). A red arrow marks a SNV present in all samples (except PBL) from case T6107 (chr19:56131557), residing in a 2kb region between two Zn-finger genes. The number at the bottom indicates the months after surgery of the primary tumor for the removal of the metastasis. **E:** Color keys for the different categories. Sample information and targeted sequence data details are provided in Supplementary Table S1 and Table S8.

**Figure 3.**
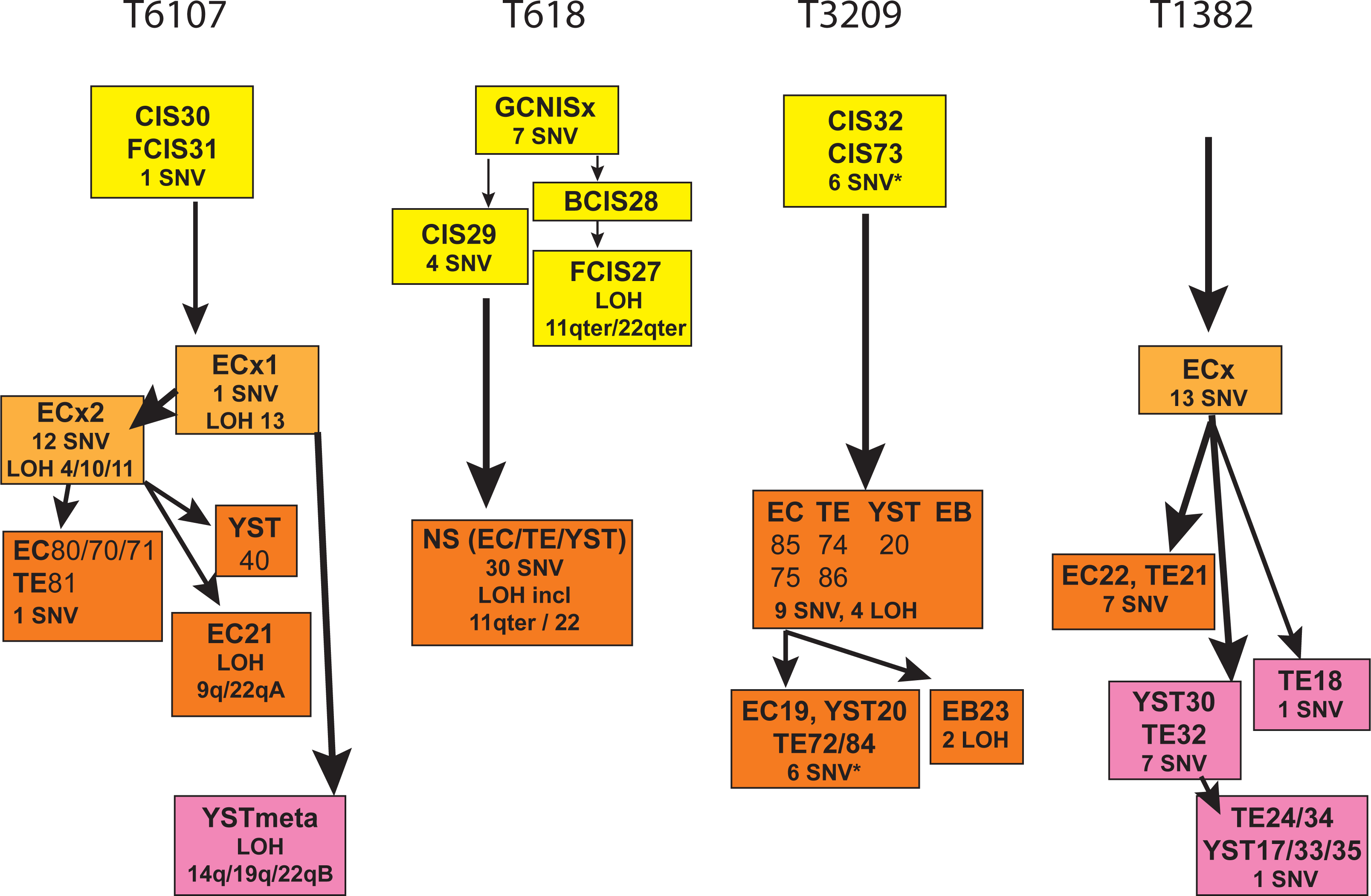
Evolution of nonseminoma. Evolutionary trees for the different histological specimens for all cases are shown on this developmental model. Coloring of boxes is according to sample type: GCNIS (yellow), precursor EC (light orange), primary tumor (dark orange) and metastasis (purple). The order and grouping of the samples is based on the similarities in allele and SNV profiles (as shown in Fig. 2 and Supplementary Fig. S7) and supported by TargetClone (Materials and Methods section), and implementing the biological constraint for NS development that the differentiated components (TE and YST) originated from an EC-type precursor. Samples with comparable profiles are grouped together. In specific cases of samples with partially non-overlapping features (i.e. SNV and/or LOH) without an immediate precursor, a non-identified precursor (GCNISx or ECx) was inserted at the branch point in the tree (T6107, T618). Similarly, to comply with the evidence that an EC is the precursor of TE and YST, a non-identified precursor ECx was introduced (T6107, T1382). Specific gains of LOH or SNV compared to their immediate precursor are indicated. *) indicates SNV present at very low read frequencies, suggesting polyclonality for GCNIS or the presence of a minor subclone in some of the primary tumor components of T3209. For case T1382, a minimal tree is presented with ordering of samples based on SNV only due to sequencing noise.

T3209: Detection of LOH (chromosomes 4, 14, 15 & 22) was markedly improved for the enriched EC sample, with increased read frequencies for most SNV (Fig. 1E). Absence of specific SNV in this EC sample and present in low read frequencies (10-20%) in the primary tumor indicated clonal variation. Major overlap in LAF patterns (LOH on the above mentioned chromosomes) and most SNV was observed for the enriched histological components and the primary NS (Supplementary Fig. S7B). A single embryonal body (EB23) showed additional regions of LOH (involving chromosomes 1 and 5). The GCNIS preparations essentially lacked allelic imbalances and SNV. The evolutionary tree for this case suggested separate developmental lineages for EB23 and four samples of EC, TE and YST (Fig. 3). The occurrence of very low frequency SNV (<5% of the reads, details Supplementary Fig. S7B) in the GCNIS preparation (which were abundant in the histological components), suggested the presence of a minor population of further progressed GCNIS.

T618: Laser capture of the histological components in the primary NS was not successful due to the presence of excess TE of cartilage differentiation. Purification of YST cells was achieved from FFPE sections using the DEPArray and provided copy number profiles comparable to the primary NS (Supplementary Figs. S1C, S2 & S10). Abundant numbers of GCNIS in the “normal” adjacent parenchyma allowed for the preparation of these cancer stem cells (Supplementary Fig. S5). Subtypes located isolated (CIS29), basal (BCIS28) or floating (FCIS27) and probably reflecting their progression state, were obtained (17). Inspection of the LAF profiles showed increasing allelic imbalances up to LOH for chromosomes 4 and 5 in these GCNIS stages (Fig. 2C/D). In addition, alterations on chromosome arms 11q and 22qter, and 7 SNV were observed in these GCNIS. Copy number analysis of DEPArray purified GCNIS did not reveal gain of chromosome arm 12p (Supplementary Fig, S10). The evolutionary tree of this case (Fig. 3) required the insertion of an unidentified precursor (GCNISx) as the progressed state GCNIS (BCIS28 and FCIS27) lacked 4 SNV (Fig. 2D) and showed loss of alleles on chromosome arms 11q and 22qter, which were retained by the primary tumor (Supplementary Fig. S8). These data strongly suggest that the abundant basal and floating GCNIS adjacent to the tumor mass represent a precursor clone that did not progress to a full malignant state and was not the founder of the primary NS.

T1382: The enriched samples were all derived from old FFPE tissue blocks and showed more amplicon drop out and noisy LAF data (Supplementary Fig. S7D). In spite of this limitation, the enriched TE and EC histologies showed good overlap for LAF, SNV, and breakpoints with the primary NS. A set of SNV, breakpoints and regions of imbalance appeared preserved in some of the metastases, while novel alterations were also observed (details in Supplementary Fig. S7D). The evolutionary tree based primarily on the SNV required the insertion of a non-identified EC precursor to explain the differences between the primary NS components and the metastases (Fig. 3).

## Discussion

Our studies on multiple samples of four cases of therapy-resistant NS provided insight in the complexity of tumorigenesis (Fig. 4). NS carried low numbers of SNV (∼0.1 per Mb, Fig. 1B), somewhat lower than reported and comparable to some of the pediatric cancers and spermatocytic tumors (12-14, 20, 31). The total number of SNV with a predicted impact on the encoded protein (3 - 18, Table 1) was similar to other reports (11-13). Overlap of mutated genes belonging to specific pathways was not observed within this small series of TGCC cases and was limited with genes reported for primary untreated TGCC (ASAP1, BRCA1, CCKBR, CROCC, CSE1L, KAT6A, KEAP1, MTOR, MYOM3 and TAF7) (11-14, 20, 32, 33).

**Figure 4.**
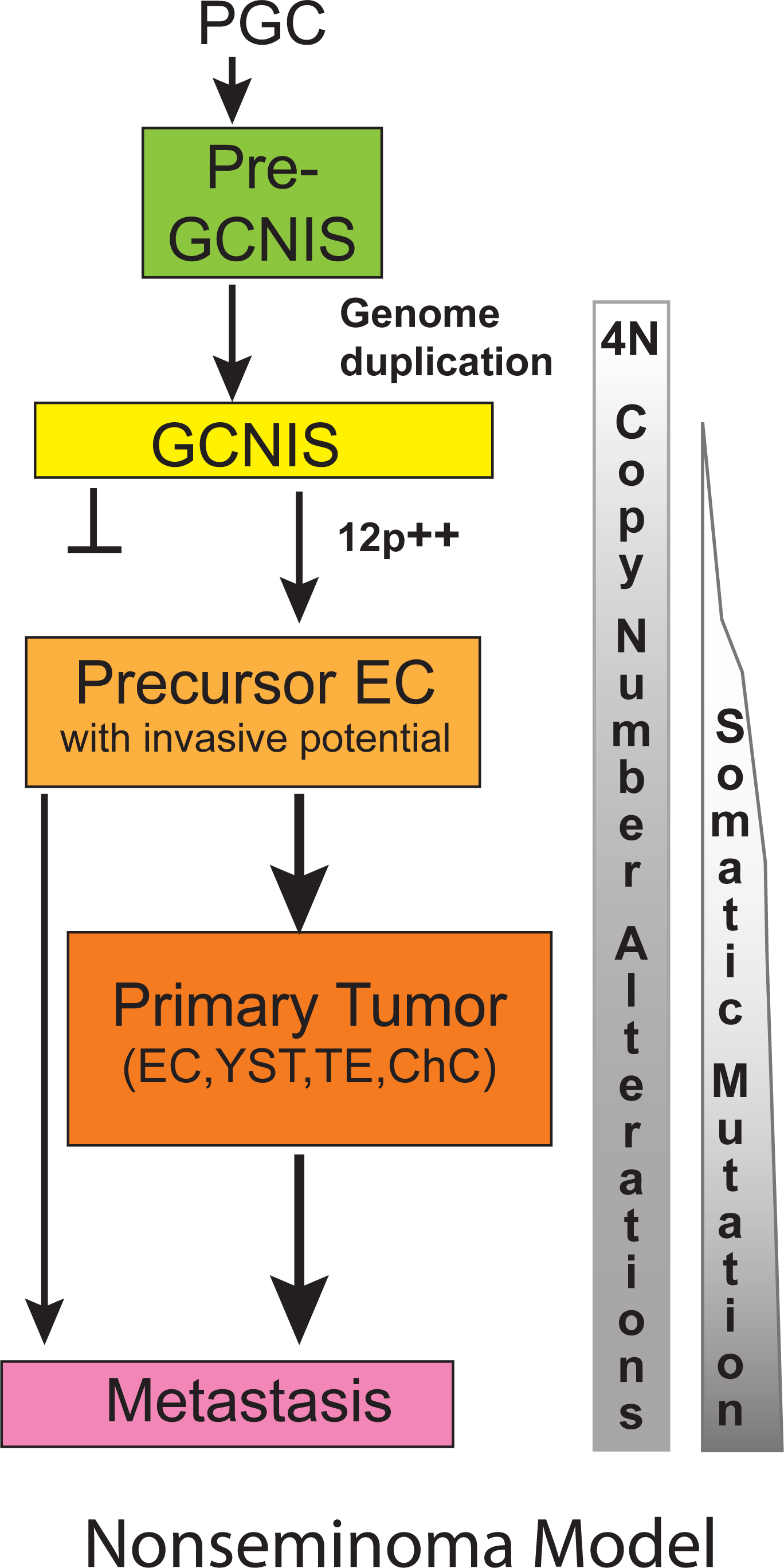
Nonseminoma evolution. A model of the different steps in NS development from normal PGC/gonocytes to metastasized NS is shown. The model is based on the available literature regarding early genome duplication, acquisition of extra copies of the short arm of chromosome 12, the pluripotent capacity of the EC (4), and the results from this study. Following initial whole genome doubling, during puberty chromosome loss may be the predominant way to change the copy numbers in the formation of GCNIS cancer stem cell (yellow). GCNIS represent a polyclonal mixture of cells, some may remain dormant and others may progress to malignancy. The gain of copies of 12p (12p^++^) is a hallmark of the precursor with invasive potential (light orange). Further losses and gains of chromosomes or chromosome fragments may contribute to the formation of the primary tumor with its distinct histological components (orange) and the typical metastases (purple). Somatic mutation appears to be limited and occurring at later stages. The metastases may also originate from precursor EC not detected in the primary tumor.

NS exhibited clonal heterogeneity which did not correlate with the histological subtypes (Fig. 3). This was expected because of EC being the stem cell component of all differentiated NS elements in the primary NS and different EC precursors providing independent lineages of differentiated cells (5, 34). A typical example is provided by the T3209 EB23 which resembles an early developing embryo derived from a single EC with a different genomic make-up (Supplementary Fig. S7B). Extensive intratumor heterogeneity was also reported for non-small-cell lung cancer (35). Our analyses also showed that metastatic clones can be derived from very early cancer stem cells that are underrepresented or even absent in the primary lesion, and not detected with the current approaches (Fig. 4). Early disseminated cells have been shown to seed metastases in models of breast, pancreatic, bladder and melanoma (36-39), but appeared less predominant in breast cancer patients (40).

The comparison of the primary tumor and the highly enriched tumor histologies revealed that the read frequencies of SNV on the autosomes (even in regions with LOH) hardly ever exceeded the 50% level (examples in Supplementary Figs. S1, S6 and S7), indicating that always a wild type copy of the particular gene was retained within the cancer genome. Similarly, the relative read frequencies of the SNV present in the GCNIS cells were low (< 0.34 for case T618, Fig. 2D) and therefore likely limited to a single allele copy. These results indicate that whole genome duplication preceded the gain of most somatic mutations (35, 40). The single SNV identified in the GCNIS of T6107 was present in high relative read frequencies (up to 50%), suggesting that this mutational event preceded genome duplication (Fig. 2B). Alternatively, this specific variant may represent mosaicism resulting from somatic mutation in early embryonic cells (41, 42). Genome doubling may also underlie development of esophageal cancer following early TP53 inactivation (43). The status of the overrepresentation of the 12p region in the purified cancer stem cells GCNIS remains uncertain, but the observed balanced allele frequencies are in line with absence of 12p gain (Fig. 2A&2C & Supplementary Fig. S8). Furthermore, LowPass WGS on DEPArray purified T618 GCNIS, revealed no gain of 12p and absence of the majority of CNA present in the primary tumor (Supplementary Fig. S10). These results are in agreement with the notion that accumulation of chr12p copies coincides with acquirement of invasive behavior (Fig. 4) (17, 44, 45).

Our data and the lack of recurrent driver mutations support the hypothesis that whole genome duplication is the primary event in NS development (Fig. 4), to be followed by overall net chromosome copy losses (4, 46). Proof for early whole genome duplication may be obtained using digital NGS based on Barcode-In-Genome technology on many individual GCNIS to determine actual chromosome copy numbers (47). Subsequently, gain of 12p copies (which may be dynamic in subclones, details Supplementary Fig. S7), gain of limited numbers of somatic mutations, and additional CNA will trigger the development of the primary NS (Fig. 4). This model is in agreement with the conclusions of Shen et al (20) for the majority of the TGCTs. Only a fraction of *KIT*-mutated SE may have acquired the mutation before whole genome duplication. In order to prevent accumulation of genomic mutations in germ cells, active surveillance and removal of PGCs with an aberrant genome is very efficient (48, 49). All PGCs are completely de-methylated and considered to be prone to aneuploidy (50), but incidence of TGCC is low in the male population. Removal of aberrant GCNIS may require functional TP53 which could be interrupted by gene mutation or by defined miRNAs (51, 52). In agreement with their extreme sensitivity to cisplatin-based therapies, TP53 mutations are extremely rare in primary TGCC indicating no selective pressure (10, 14, 52). The presence of an amplified MDM2 locus in two metastases of case T6107 (Supplementary Fig. S9) may have provided an alternative route for inactivation of TP53 and therapy resistance (10, 32, 53). The BRCA-like base substitution signature of this cancer hints at inefficient homologous recombination repair (54). This BRCA-like signature was not reported in a recent exome sequencing of TGCC (20). There is currently no evidence for the direct involvement of the *BRCA* genes (i.e. absent pathogenic gene mutation and no rearrangement signature (30), and no association of TGCC with familiar *BRCA1/2* mutations), although increased methylation of the *BRCA1* gene promoter was observed in some NS without EC (20). It is tempting to speculate that other components of this homologous recombination repair pathway may be affected and responsible for the specific base substitution signature in the absence of direct involvement of BRCA genes (30, 54). Alternatively, BRCA-related repair functions may be low or turned off intrinsically in the embryonic PGC/gonocytes (55) and during early TGCC development, and thus contribute to the accumulation of this specific pattern of base substitutions. Therefore, TGCC development may be the result of the properties of the PGC (de-methylated and reduced repair pathway), which allows the tetraploid GCNIS to evade apoptosis and survive till puberty and subsequently progress to malignancy due to genomic destabilization (56). The (epi)-genetic triggers for destabilization of the tetraploid genome and initiation of the development of malignant clones remain as yet largely unknown, although the induction of mitogenic signaling by testosterone may contribute.

The observed heterogeneity in the primary tumor, metastases and precursor lesions of nonseminoma may impact on clinical decisions and treatment strategies. The occurrence of metastatic tumors with little molecular overlap with the primary lesion indicates that treatment of therapy-resistant recurrences should be targeted at their molecular properties according to the concepts of personalized medicine. The identification of abundant cancer stem cells GCNIS which did not contribute to the development of the NS further complicates the clinical advice to patients with GCNIS only disease (57). In view of its tendency to progress in 70% of the patients within 7 years, novel predictive markers for the GCNIS progression are needed. The BRCA-like mutational signature in TGCC indicative for reduced homologous recombination repair, may be suggestive for combined use of PARP inhibitors and platinum-based therapy (20, 58) but requires additional support. Further studies into NS and SE cases, with GCNIS adjacent to the primary tumor and different histologies or localization, are needed to identify reliable markers for GCNIS progression, to unravel the critical steps for malignancy and therapy resistance, and to decipher the origin of the BRCA-like mutational signature of TGCC.

### Additional Information

#### Ethics approval

Use of tissue samples remaining after diagnosis for scientific reasons was approved by Medical Ethical Committee of the Erasmus MC Rotterdam (The Netherlands), permission 02.981. This included the permission to use the secondary tissue without further consent. Samples were used according to the “Code for Proper Secondary Use of Human Tissue in The Netherlands” developed by the Dutch Federation of Medical Scientific Societies.

#### Availability of data and materials

WGS, ENA PRJEB20644, accession numbers ERX2100523 – 530; targeted sequencing, ENA PRJEB20644, accession numbers: ERX2019898-958; RNA-seq, ArrayExpress, accession number: E-MTAB-5746; DNA methylation, GEO (GSE58538, GSM1413103 to GSM1413106) or ArrayExpress (E-MTAB-5842, sample T6107-YSTmeta).

#### Conflict of interest

The authors declare no potential conflicts of interest.

## Funding

Financial support: Whole genome sequencing was provided by Complete Genomics, Inc. Mountain View, CA 94043, USA, and DEPArray experiments were performed by Menarini Silicon Biosystems S.p.a. (Castel Maggiore, Italy).

## Authors’ contribution

LD and LL: design of experiments and writing the manuscript. AG, HS, RM, LD: execution of experiments, JvR, HW, MN, JdR & LD for bioinformatics analyses, JWO: pathology review. All authors approved the manuscript.

## Acknowledgements

We greatly acknowledge Lieke Dons and Dragana Bertovic for assistance in sample purification. We thank Remko Hersmus for performing FISH experiments; Daphne Heijsman, Arne S. IJpma and Peter van de Spek for help with WGS data; Saskia D. Hiltemann, and Andrew P. Stubbs for support with Galaxy and analysis of structural variants (IFuse); Ans van den Ouweland for functional evaluation of SNV; Erik-Jan Dubbink for targeted sequencing options, and Alex Nigg for the plug-in to enhance FISH images. We thank the microarray unit of the DKFZ Genomics and Proteomics Core Facility for providing the Illumina Human Methylation arrays and related services, and Volker Hovestadt and colleagues for help with the copy number analyses of methylation intensity data. We also acknowledge the Erasmus MC Cancer Computational Biology Center for giving access to their IT-infrastructure and software that was used for computations and/or data analysis in this study. This study was supported in part by a WGS grant from Complete Genomics, Inc. Mountain View, CA 94043, USA. We appreciate the support from Rosanna Lanzellotto, Alberto Ferrarini, Bechir Boughaba, Raimo Tanzi and Francesca Fontana from Menarini Silicon Biosystems S.p.a. Castel Maggiore (Italy) for the DEPArray experiments.

Supplementary information is available at the British Journal of Cancer’s website

